# Sodium Diethyldithiocarbamate antiparasitic activity against different *Trypanosoma cruzi* strains: Insights of its biological activity

**DOI:** 10.1101/2020.07.06.189233

**Authors:** Johny Wysllas de Freitas Oliveira, Taffarel Melo Torres, Cláudia Jassica Gonçalves Moreno, Bruno Amorim-Carmo, Igor Zumba Damasceno, Ana Katarina Menezes Cruz Soares, Jefferson da Silva Barbosa, Hugo Alexandre Oliveira Rocha, Marcelo Sousa Silva

**Affiliations:** Immunoparasitology Laboratory, Department of Clinical and Toxicological Analysis, Centre of Health Sciences, Federal University of Rio Grande do Norte, Natal, Brazil; Programa de Pós-graduação em Bioquímica, Centro de Biociências, Universidade Federal do Rio Grande do Norte. Natal, Brazil; Centro de Ciências Biológicas e da Saúde- Universidade Federal Rural de Semiárido. Mossoró, Brazil; Programa de Pós-graduação em Ciências Farmacêuticas, Centro de Ciências da Saúde, Universidade Federal do Rio Grande do Norte. Natal, Brazil; Departamento de Engenharia de Materiais, Centro de Tecnologia, Universidade Federal do Rio Grande do Norte. Natal, Brazil; Laboratório de Biotecnologia de Polímeros Naturais – BIOPOL, Departamento de Bioquímica, Centro de Biociências, Universidade Federal do Rio Grande do Norte. Natal, Brazil; Instituto Federal de Educação, Ciência e Tecnologia do Rio Grande do Norte (IFRN) - *Campus* São Gonçalo do Amarante. São Gonçalo do Amarante, Brazil; Global Health and Tropical Medicine, Instituto de Higiene e Medicina Tropical, Universidade Nova de Lisboa. Lisbon, Portugal

**Keywords:** Chagas disease, *Trypanosoma cruzi*, Sodium Dietyldithiocarbamate, Antiparasitic activity, DTU’s, Neglected tropical disease, drug discovery

## Abstract

**Background:** Chagas disease is caused by the protozoan *Trypanosoma cruzi*, a neglected tropical disease that affects thousands of people, mainly in Latin America. The drugs currently used in therapy are toxic and have therapeutic limitations during treatment. In addition, the genetic diversity of *T. cruzi* represents an important variable and challenge with regard to the pathogenesis of the infection, the epidemiological profile of the cases, and the therapeutic control of the infection. Sodium diethyldithiocarbamate (DETC) is a compound of high pharmacological versatility acting as metal chelators and producing reactive oxygen species. Thus, the objective of this work is to characterize the antiparasitic action of DETC against different strains and evolutionary forms of *T. cruzi*, as well as the characterization of the mechanism of antiparasitic action.

**Methodology/Principal findings:** The different strains and evolutionary forms of *T. cruzi* were grown in LIT medium. To evaluate the antiparasitic activity of DETC, the evolutionary forms epimastigote and trypomastigote of *T. cruzi* were used by resazurin reduction methods and by counting under optical microscopy. Different response patterns were obtained between the strains and an IC_50_ of DETC ranging from 9.44 ± 3,181µM to 60.49 ± 7.62 µM. Cell cytotoxicity against cell lines 3T3 and RAW and evaluated by MTT, demonstrated that DETC in high concentration (2222 µM) reduces around 60% the cell capacity of MTT reduction. The antiparasitic activity of DETC has been demonstrated through damage caused in the mitochondria of *T. cruzi*, a reduction of up to 80% in the mitochondrial potential of the parasites, as well as through damage caused in the membrane of the parasite.

**Conclusion:** In this study we can conclude that DETC has antiparasitic activity against different genotypes and evolutionary forms of *T. cruzi*, representing a promising molecule as a drug for the treatment of Chagas disease.

## 1. Introduction

Chagas Disease (CD) is a Neglected Tropical Disease (NTD) caused by flagellated protozoan *Trypanosoma cruzi*. This disease affects more than 8 million people worldwide, causing more than 10,000 deaths per year and has more than 80 million people living in risk zone [1]. Chagas disease has an acute asymptomatic phase or nonspecific clinical signs, and in the chronic phase, individuals may be asymptomatic or progress to cardiac and/or digestive complications [2]. This disease can be transmitted in different ways and the two main transmission mechanisms are vector transmission, during the blood meal of triatomines, and ingestion of food contaminated with the faeces of these vectors [3]. However, other transmission mechanisms also contribute to Chagas disease, such as blood transfusion, organ transplants, and vertical congenital transmission responsible for the dissemination of disease in other countries through the emigration for non-endemic countries [4].

*T. cruzi* is a flagellated protozoan belonging to the trypanosomatid family. This parasite has a complex heteroxenic life cycle, maintaining its biological cycle between the invertebrate host of the *Triatomine* family and the mammalian host (humans, dogs, and wild animals) passing by metacyclogenesis during this cycle suffering alteration in evolutionary form [5]. Although *T. cruzi* is represented by a single species, the diversity and genetic variability of these parasites are made up of different strains grouped based on their genetic characteristics named Discrete Typing Units (DTUs) [6]. Thus, *T. cruzi* can be classified into seven different genetic variants or genotypes in which was included several strains, called from TcI to TcVI, and, more recently, the variant called TcBat [6-7]. These genetic variants of *T. cruzi* are important from an epidemiological point of view, as they are involved in different aspects of Chagas disease, mainly in the context of the transmission cycle, clinical manifestations, pathogenesis, reservoir and geographical distribution of these different strains of *T. cruzi* [8].

In the context of pharmacological therapy for Chagas disease, the drugs currently used are benznidazole and nifurtimox [9]. These two nitroaromatic-compounds act on the parasite by producing reactive oxygen species (ROS), thus presenting antiparasitic activity. However, from a clinical point of view, these drugs have limitations regarding their use, since they are responsible for several adverse reactions like liver damage, pruritus, spots on the skin, and itchy in eyes [10]. Additionally, these drugs have low pharmacological efficacy in the chronic phase of infection with *T. cruzi* [11]. This fact has led to the search for new compounds that may be more effective than these drugs in the treatment of Chagas disease.

Diethyldithiocarbamate (DETC) is a compound belonging to the class of dithiocarbamates formed by two ethyl substituents linked to an amine group, which in turn is linked to carbon disulfide. The chemical structure of DETC has two distinct portions; the first, an amine portion formed by nitrogen and the two ethyl groups, which stimulate the production of reactive oxygen species. A second portion, a carbon disulfide end that can chelate on metals, thus attributing an essential biological activity of this compound against metalloproteases, enzymes necessary for parasitic biology [12-14]. Due to the structural characteristics of DETC, several pharmacological applications were observed in this compound [15-21]. In addition, we recently published a review that addresses the different justifications for enhancing DETC as a potential antiparasitic drug [15].

However, little is known about the antiparasitic activity of DETC against *T. cruzi*. Thus, this study’s objective is to evaluate the antiparasitic activity of DETC *in vitro* gainst the different strains (different DTUs) and evolutionary forms of *T. cruzi*. Thus, in this work we report for the first time the antiparasitic activity of DETC against different strains of *T. cruzi*, as well as insights of its mechanisms of action as antiparasitic drug.

## 2. Methods

### 2.1. Axenic cultivation of *Trypanosoma cruzi*

Different strains of *T. cruzi* TcI (strain Dm28c) [22]; TcII (strain Y) [23]; Tc (QMM9 strain) [24] and TcVI (CLBlener strain) [25] were grown in LIT (Liver Infusion Tryptose) medium supplemented with 10% inactivated fetal bovine serum (FBS) (v/v) and 5% of antibiotic streptomycin/penicillin (100 UI/mL). All strains were donated by *Laboratório de Parasitologia* (FCFAr), Araraquara, Brazil. The epimastigote forms of *T. cruzi* were kept at 27 °C in a BOD oven (incubator chamber, ASP, SP-500). The trypomastigote forms of *T. cruzi* were obtained through the nutritional stress method, from epimastigote forms of the parasite for a period of 25 days, according to the methodology developed by Camargo et al. [26]. The morphological changes were confirmed by microscopy considering the cultures that presented in their parasitic concentration higher than 75% of the trypomastigote forms.

### 2.2. Cells

RAW 264.7 macrophage cell line (ATCC® number TIB-71™) and 3T3 fibroblast (ATCC® CRL-1658™) was cultured in DMEM (Cultilab, Campinas, SP, Brazil) supplemented with 10% (v/v) fetal serum bovine (FBS) (Cultilab, Campinas, SP, Brazil) and antibiotics (100 U/mL penicillin and 100 µg/mL streptomycin). The cells were incubated at 37 °C in a humidified atmosphere with 5% CO_2_. For maintenance of the cells, the culture medium was changed every three days, and the cells were further subcultured at the 80% confluence using a cell scraper (RAW cells) or trypsin/EDTA (3T3 cells).

### 2.3. Determination of DETC antiparasitic activity against different strains and evolutionary forms of *Trypanosoma cruzi*

The different strains and evolutionary forms of *T. cruzi*, mentioned above, were used in this study. The parasites were diluted to a concentration of 1.0 × 10^7^ parasites/mL and then kept 200 µL in 96-well plates together with different concentrations of sodium diethyldithiocarbamate (DETC), ranged from 4.44 to 444.00 µM. The antiparasitic activity of DETC was evaluated through the viability of the parasites after a period of 24 and 48 hours of cultivation. As a positive control of antiparasitic activity against *T. cruzi*, benznidazole (BZN) was used in different concentrations, ranged from 3.84 to 500.00 µM, this range was selected due to existing variability response against the different strains of *T. cruzi* and to determinate the IC_50_ of BZN to compare with DETC [7].

Parasite viability was determined by the resazurin (Sigma Aldrich, Laramie, WY, USA) reduction assay at a concentration of 1 mM and subsequently measured by spectrophotometer (Epoch, BioTek Instruments, Winooski, VT, USA), at 570nm and 600nm wavelengths [27]. The antiparasitic activity of DETC was expressed as % of resazurin reduction = 100 – [(Atest_570_ – (Atest_600_*Ro))/(Acontrol_570_ - (Acontrol_600_*Ro)] × 100, in which Atest corresponds to the absorbance of the experimental group and Acontrol corresponds to the absorbance of the negative control, 570 and 600 are the wavelengths to corresponding 570 nm and 600 nm and Ro represent the index of correction between medium and resazurin. Based on % resazurin reduction was calculated the IC_50_ that represents the drug concentration to reduce by fifty percent the parasite population.

The viability of the parasites was also determined by microscopy in a clear camera through direct counting in a Neubauer camera. In this essay, parasites with some movement were considered viable. Cell viability results were compared between the two methodologies. In some conditions, when DETC promoted 100% mortality, the culture the medium containing the parasites was collected from each plate, centrifuged (2000 rpm, 10 min, 4 °C) to remove the DETC and the old medium. Afterwards, the parasites received a LIT medium, supplemented with 10% (v/v) FBS and 5% of antibiotic streptomycin/penicillin (100 UI/mL). After 7 days, the culture was analysed by conventional optical microscopy. The absence of live parasites confirms 100% mortality.

### 2.4. Evaluation of membrane structure alteration in *Trypanosoma cruzi* submitted to DETC by scanning electron microscopy

To evaluate possible morphological changes in the membrane of different strains of *T. cruzi* caused by DETC, the parasite membrane structure was analysed by scanning electron microscopy after treatment with DETC, according to a protocol established by Amorim-Carmo and colleagues [30]. Briefly, the epimastigote forms of the different strains of *T. cruzi* (1.0 × 10^7^ parasites/mL), in the same cultivation conditions described previously in topic 2.2, were treated with DETC (44.40 µM and 222.00 µM) in identical conditions described previously. After 24 hours, the parasites were centrifuged for 10 min at 2000 rpm (4 °C) and washed twice in PBS 1x pH 7.4. The parasites were then fixed in 2.5% (v/v) PBS-glutaraldehyde for 4 hours. After this step, the parasites were dehydrated being exposed to different ethanol concentrations. First, to dehydrated the parasite was added ethanol at 25% and leave for 10 minutes, following by washes in PBS 1x pH 7.4 for 10 minutes at 1500 rpm room temperature, the same operation was repeated in 50%, 80%, and 100% ethanol concentration to dehydrate the parasite for next steps. Then, the parasites were resuspended in absolute ethanol, was dropped in silicon plates and placed to dry at room temperature. Finally, after dried, the parasites were placed on stubs and subjected to metallization with gold using the sputtering (Bal-Tec SCD-005 Sputter Coater, Schalksmühle, NWR, GER) in an argon atmosphere for 30 seconds with a current of 30 mA and then analysed by scanning electron microscopy under a FEG microscope (Model augira, Brand Carl Zeiss, Oberkochen, WB, GER).

### 2.5. 3-(4,5-dimethylthiazol-2-yl)-2,5-diphenyltetrazolium bromide) tetrazolium (MTT) reduction assay on cell viability submitted to DETC

The ability of RAW 264.7 macrophages and fibroblast 3T3 to reduce MTT was evaluated according to the previously described method of Mosmann [28]. Initially, the cells were seeded in 96-well plates at a density of 5 × 10^3^/well and kept at culture condition for 12 hours. After, the medium was replaced by a medium contained DETC at the different concentrations tested (from 4.44 to 2,222 µM). After 24 h, the culture medium was replaced with 100 µL of MTT (1 mg/mL dissolved in DMEM). Then, the cells were incubated for 4 h at 37 °C. Subsequently, the culture supernatant was discarded, and the crystals of formazan were solubilized in ethanol, 100 µL/well. Absorbance was measured with an Epoch microplate spectrophotometer (Biotek Instruments Inc., Winooski, VT, USA) at 570 nm. Cell viability was calculated in relation to the negative control using the formula: % viability = (Atest/AControl) × 100, in which Atest corresponds to the absorbance of the experimental group and Acontrol corresponds to the absorbance of the negative control. Based on % MTT reduction was calculate the IC_50_ that represents the drug concentration to reduce by fifty percent the cell population.

### 2.6. Determination of Selective Index of DETC

To assess of predilection of DETC between the parasite and cell was used the Selective Index (SI). This Index represents the preference of drug calculated by: SI = IC_50parasite_/IC_50cell_, in which IC_50parasite_ represents the drug concentration to reduce by fifty percent the parasite population in *vitro* and IC_50cell_ represents the drug concentration to reduce by fifty percent the *in vitro* cell population.

### 2.7. Evaluation of death mechanism in *Trypanosoma cruzi* caused by DETC

Annexin V/propidium iodide (PI) was used to evaluate the DETC death mechanism at *T. cruzi*. Briefly, the epimastigotes and trypomastigotes forms of the different strains of *T. cruzi*, in the same culture conditions described previously in topic 2.2, were exposed to DETC (4.44 µM, 44.40 µM or 444.00 µM) for 24 hours. Then, the parasites were centrifuged (2000 rpm for 5 minutes, 4 °C), rinsed with PBS 1x pH 7.4 (Phosphate-Saline Buffer), and suspended in 300 µL Bind Buffer 1x. Then, 5 µL of Annexin V-FITC (fluorescein isothiocyanate) was added to the system, let reacting for 10 minutes. Then, the parasites were centrifuged (2000 rpm for 5 minutes, 4 °C) and resuspended again in 200 µL Bind Buffer 1x and 10 µL of PI. All these steps are described in the protocol proposed by the manufacturer (Annexin V FITC Apoptosis Detection Kit, Invitrogen, Carlsbad, CA, USA). After 10 minutes, the fluorescence intensity was determined using a flow cytometer (FACSCanto II, BD Biosciences, Eugene, OR, USA) with FACSDiva software, version 6.1.2 (Becton Dickson, Franklin Lakes, NJ, USA). For the analysis, the 10,000 events count determined for each evolutionary form and strain of *T. cruzi* was used.

### 2.8. Evaluation of mitochondrial damage in *Trypanosoma cruzi* by DETC

To analyse the mitochondrial damage induced in different strains and evolutionary forms of *T. cruzi* by DETC, the mitochondrial potential marker rhodamine-123 (Invitrogen, Carlsbad, CA, USA) was used, following the manufacturer’s recommendations. Briefly, the different strains and evolutionary forms of *T. cruzi* were treated with different concentrations of DETC (44.40 µM; 111.00 µM; or 222.00 µM) for 24 hours in same conditions described previously. After treatment, the parasites were centrifuged at 2000 rpm for 6 min at 4 °C and rinsed with PBS 1x pH 7.4. Then, the parasites were suspended and 200 µL of PBS and 0.5 µL of rhodamine-123 (5 mg/mL) were added, as described by Sulsen and colleagues [29]. After 30 minutes, the parasites were rinsed twice in PBS and analysed by flow cytometer at wavelengths of 488 and 633 nm Finally, the fluorescence intensity was determined by flow cytometer (FACSCanto II, BD Biosciences, Eugene, OR, USA) with FACSDiva software, version 6.1.2 (Becton Dickson, Franklin Lakes, NJ, USA). The mitochondrial activity of the parasites was calculated according to the formula: % of reduction rodamine stain = (fluorescence intensity of the parasites treated with DETC)/(fluorescence intensity of negative control) * 100. A total of 10,000 events were analysed for each strain and evolutionary form of *T. cruzi*.

### 2.9. Statistical analysis of the data

All experiments were carried out in triplicates and independently, the results are present in mean ± standard deviation. The data were submitted to the Shapiro-Wilk normality test. Parametric data were analysed using the ANOVA test associated with the Tukey-T post-test and Pearson correlation. Tests were performed using the software GraphPad Prism v. 7.0 (2016) and P.A.S.T v. 2.17 (2012).

## 3. Results

The parasite’s ability to reduce resazurin after exposure to different conditions is shown in Figure 1. In general, this parasite’s ability was affected depending on the time (it was most affected after 48 h) and the concentration of DETC. In addition, the effect of DETC also varied according to each strain on different evolutionary forms; epimastigotes forms were more affected than trypomastigotes forms.

**Figure 1:**
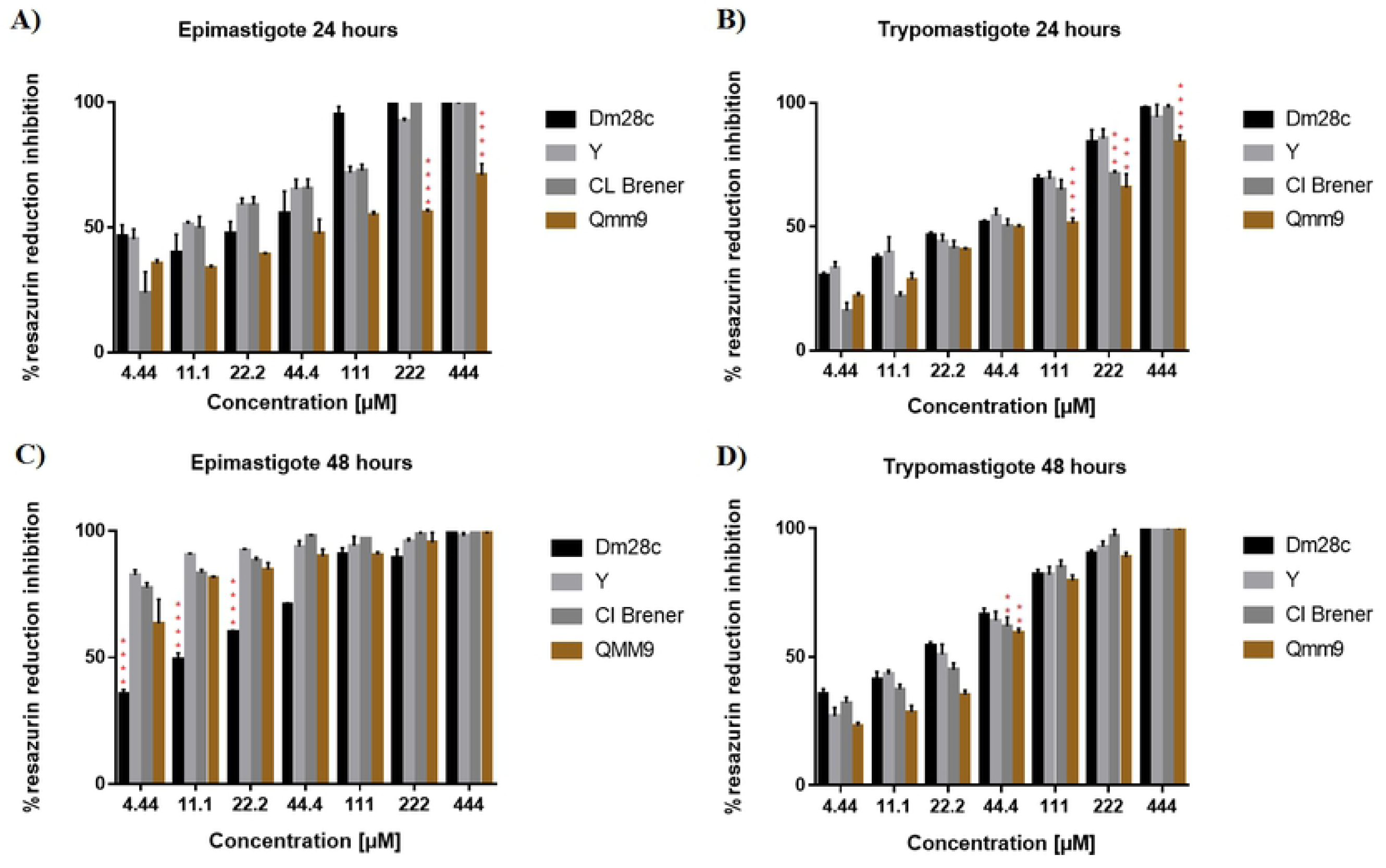
Evaluation of resazurin reduction inhibition by different strains and evolutionary forms of *Trypanosoma cruzi* treated with DETC. Parasitic inhibition of resazurin reduction (%) of DETC against different forms (epimastigotes and trypomastigotes) and strains (Y, Dm28c, Cl Brener, and Qmm9) of *T. cruzi*, after 24 and 48 hours of culture and analysed by Resazurin assay. A: Epimastigotes for 24 hours of treatment; B: Trypomastigotes for 24 hours of treatment; C: Epimastigotes for 48 hours of treatment; D: Trypomastigotes for 48 hours of treatment. Results presented as mean ± standard deviation of the percentage of parasitic inhibition in a triplicate system and for the statistical analysis of the Anova Test together with the Tukey Post-test (P <0.01 (**); P <0.001 (***), P<0.0001(****). In statistical analysis was compared the response in different strains submitted to the same concentration of DETC.

The strain Y was the most sensible to exposure at DETC, suffering a significant reduction of resazurin reduction capacity. QMM9 in both forms, epimastigote and trypomastigote of all strains, was less affect to the ability to reduce resazurin after 24 hours of exposure at DETC. Among all conditions tested with DM28c, the one in which the greatest resistance of the parasites was observed when the DB28c epimastigote forms were exposed for 24 h to the DETC. Interestingly, after 48 hours of exposure at compound, all strains tested in both evolutionary forms had the capacity to reduce resazurin virtually reduced to zero.

To confirm the DETC toxicity against parasite, they were also counted in light chamber, which was considered viable just parasites present movement. The results of counting are present in Figure 1S. When compared the results present in Figure 1 and Figure 1S, they present similarities when associating the capacity of reducing resazurin and movement of the parasite, both results to all strains present similar profile of answer at DETC.

The IC_50_ of DETC was determined using the data from resazurin assay and the values are showed in Table 1. The IC_50_ was different from each other in most of conditions tested and the IC_50_ of BZN, in the same conditions of DETC, was always higher when compared to IC_50_ of DETC for all strains in each evolutionary form.

**Table 1:** DETC antiparasitic activity, expressed in IC_50_ values ± standard deviation, against the different strains and evolutionary forms of *Trypanosoma cruzi* during 24 hours of cultivation.

| *Trypanosoma cruzi*<br>strains (DTUs) | IC<sub>50</sub> epimastigote |  | IC<sub>50</sub> trypomastigote |  |
| --- | --- | --- | --- | --- |
|  | DETC (µM) | BZN (µM) | DETC (µM) | BZN (µM) |
| Strain Dm28c (TcI) | 15.94±7.43 | 69.32±8.42 | 25.00±5.34 | 89.95±4.87 |
| Strain Y (TcII) | 9.44±3.181 | 92.66±7.78 | 23.35±5.41 | 98.74±3.98 |
| Strain QMM9 (TcIII) | 60.49±7.62 | 99.38±9.41 | 53.49±8.47 | 108.71±6.74 |
| Strain CI Brener (TcVI) | 15.18±3.64 | 62.86±8.47 | 43.18±7.61 | 82.43±5.83 |

BZN - benznidazole

The strain QMM9 was less affected, by exposure to DETC, in the capacity of reducing resazurin since it showed the highest IC_50_ between tested strains. Meanwhile, with the smaller IC_50_, the strain Y was the most affected by the exposure to DETC. The strains Dm28c and Cl Brener present a similar profile in epimastigote form, but different answer profile in trypomastigote form. Furthermore, the strains Dm28c, Y and Cl Brener present a higher IC_50_ in trypomastigote form when compared with the result in epimastigote form. At the same time, QMM9 had a reduced in IC_50_ of epimastigote form compared with the trypomastigote form.

In addition, the correlation between each IC_50_ result of DETC and BZN against all strains in different evolutionary forms was also determined. Based on results, it was observed through the Pearson Correlation a R square > 0.88 to all strains tested, representing a strong correlation between results of DETC and benznidazole, in other words when DETC had lesser efficacy the same occurred with BZN and the opposite can apply either.

In addition, as in some conditions, DETC was able to reduce in 100% of capacity to reduce resazurin by different strains and confirm the relation of 100% of capacity to reduce resazurin with 100% of mortality, it was made the re-cultivation. For that, the parasites were submitted to the same treatment described above, and after the treatment period, the parasites were placed in another plate containing the growth medium without the tested compounds. After seven days, the recipient was analysed by optical microscopy and the presence of viable parasites was not detected.

To observe if the exposure to DETC by different strains of *T. cruzi* causes morphological changes in membrane structure of parasite, after treatment the parasites had their membrane structure analysed by scanning electron microscopy (SEM) and results are shown in Figure 2. To this essay it was utilized just the strain Y, the reasons for that is due be the most susceptible strain, when analyse the results in resazurin reduction in Figure 1, making more evident the finds in SEM. In Figure 2.A is shown the structure of parasite without any treatment, it presents a smooth membrane, without the presence of roughness or filaments, in addition, it presents the elongation of the flagellum. In Figure 2.B it is possible to observe pores and the presence of slight roughness in the parasite membrane when the parasite was exposed to 44.4 µM of DETC. In Figure 2.C and2.D, when parasite was exposed to 222.0 µM, it is possible to observe several pore formations in membrane structure of parasite. In Figure 2.C a rosette structure is presents, common to *T. cruzi* culture, showing lot of pores in several parasites. Specifically in Figure 2.D, excessive roughness is observed in the parasite membrane of and loss of conformation. The morphological alterations present in strain Y, after treatment with DETC, should also occur in other strains.

**Figure 2:**
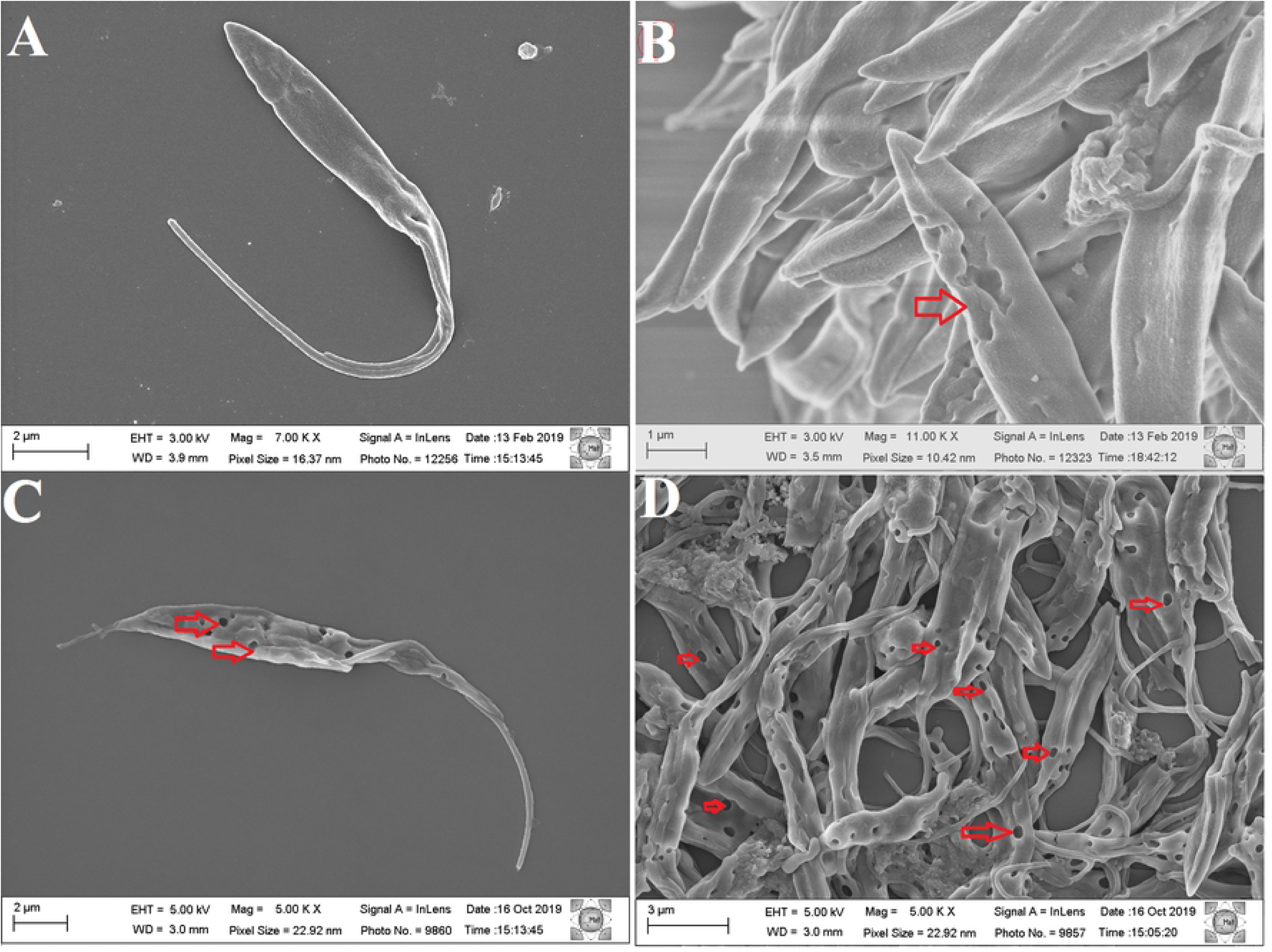
Morphological changes of *Trypanosoma cruzi* epimastigotes treated with diethyldithiocarbamate and analysed by scanning electron microscopy. A: Untreated parasite; B: Parasites treated with 44.4 µM DETC; C and D: Parasites treated with 222.0 µM DETC; the arrows indicate the presence of pore in the parasite membrane structure.

The next step was to assess the DETC effect on the ability of mammalian cells to reduce MTT. Therefore, RAW and 3T3 cells were submitted to DETC (from 4.44 to 2222.0 µM) for 24 hours, as described in methods section. As shown in Figure 3, DETC affected the cells’ ability to reduce MTT. Low concentrations of DETC affected RAW cells more than 3T3 cells. From the concentration of 11.1 µM to 111.0 µM, the cell line RAW was more sensitive, statically observed from concentration 222.0 µM, RAW and 3T3 had similar profiles of response against DETC. In this case, the ability of both cells to reduce MTT decreased to around 50%, and did not decrease even with the increase in the concentration (from 1111.0 to 2222. 0 µM) of DETC.

**Figure 3:**
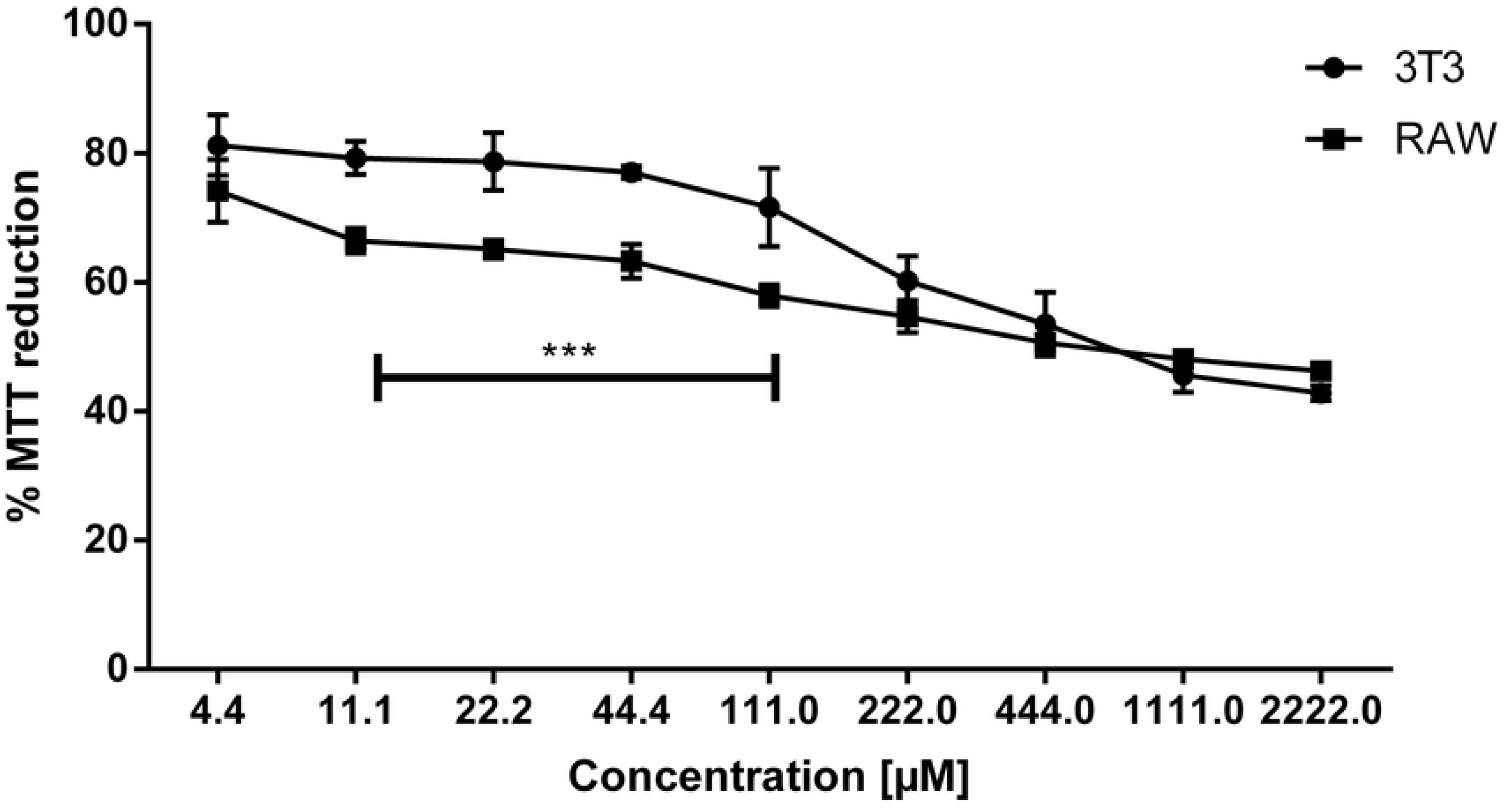
Evaluation of the MTT reduction in 3T3 and RAW cell cultures exposed to diethyldithiocarbamate. Viability curve of the 3T3 and RAW cell lines obtained through the MTT reduction assay and cells treated with different concentrations of DETC for 24 hours, 100 % represent cells untreated. Results presented as mean ± standard deviation of the percentage of MTT reduction for each cell line in a triplicate system and for the statistical analysis the Anova Test, together with the Tukey Post-test, were used to compare the reaction of two cell lines against DETC treatment, showing statistical difference in response of range concentration from 11.1 µM to 44.4 µM P <0.001 (***).

The IC_50_ of each cell line was determinate based on the decrease of reduce MTT when expose to DETC. Then, based on the MTT metabolization curve present in Figure 2, for each cell line, the IC_50_ values 859.90 µM and 698.80 µM, for cellular lineages, 3T3 and RAW, respectively, was calculated.

The selectivity index (SI) was determined based on the data from the resazurin and the MTT assays, as described in the Methods section. Table 2 shows the different IS of DETC obtained with different strains and evolutionary forms of *T. cruzi*. In addition, the different strains on different evolutionary forms associated with each cell line presented different SI. The selectivity of DETC against strain QMM9 towards the cell lines 3T3 and RAW presented the smaller values of SI for both evolutionary forms and cellular lineages. Meanwhile, selectivity against strain Y towards cell lines presented higher values of SI for both evolutionary forms and cellular lineages. In addition, the SI of DETC against strains Cl Brener and Dm28c presented a similar epimastigote profile, but the SI trypomastigote profile is different, being SI of Cl Brener closer of QMM9. Already, the SI of DETC against Dm28c trypomastigote was like the strain Y.

**Table 2:** Selectivity index (SI) of diethyldithiocarbamate in different strains and evolutionary forms of *Trypanosoma cruzi* after 24 hours of cultivation.

| *Trypanosoma cruzi*<br>strains (DTUs) | 3T3: Epi | 3T3: Trypo | RAW: Epi | RAW: Trypo |
| --- | --- | --- | --- | --- |
| Strain Dm28c (**TcI**) | 53.943 |  | 43.839 |  |
| Strain Y (**TcII**) | 91.062 | 34.396 | 74.002 | 27.952 |
| Strain Qmm9 (**TcIII**) | 14.216 | 36.827 | 11.552 | 29.927 |
| Strain CI Brener (**TcVI**) | 56.647 | 16.076 | 46.034 | 13.064 |
|  |  | 19.914 |  | 16.183 |

**Note:** Evolutionary forms of *T. cruzi* epimastigote (Epi) and trypomastigotes (Trypo) and cell lines (3T3 and RAW cells).

The fluorescence profile of death cellular markers annexin/PI in different strains of *T. cruzi* when exposed to DETC is shown in Figure 4. Based on these results it is possible to observe the absence of death cellular markers annexin/PI in all strains, when exposed to different concentrations of DETC (4.44 µM, 44.4 µM, 444.0 µM). These concentrations were used based on results of resazurin and count, seeking to identify the mechanism of death related to the stain annexin/PI in flow cytometer. The data for strains Y and Dm28c when treated with DETC was like those observed with untreated parasites, that is, more 97% of parasites was considered viable even when DETC was used at higher amount (444.0 µM). However, a low fluorescence response of death cellular markers annexin/pi was verified to strains Cl Brener and QMM9, when exposed to highest concentration of DETC. 71 % and 80% of viability was identified for both strains, respectively, after exposure.

**Figure 4:**
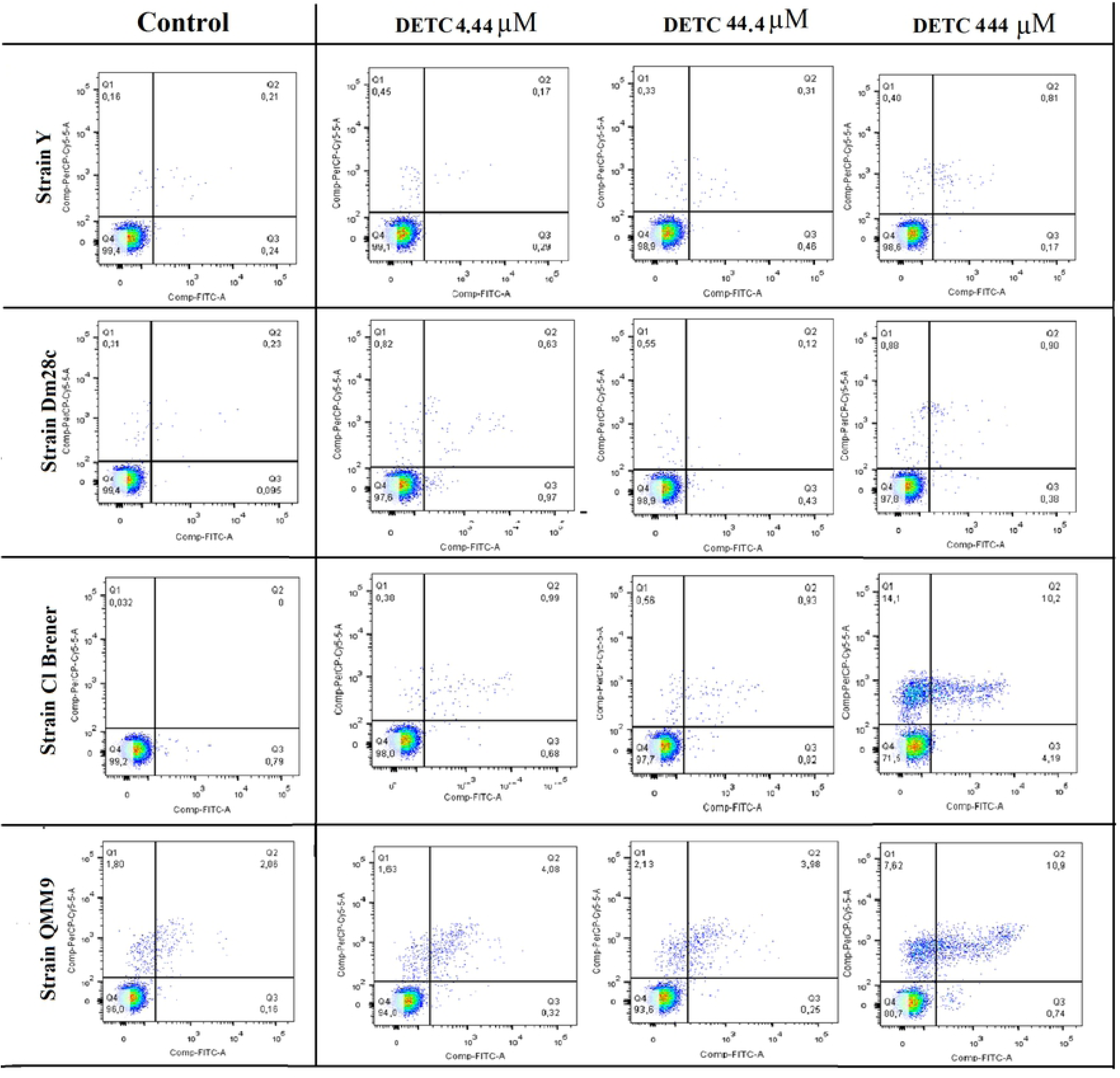
Analysis of the cell death mechanism of trypomastigotes from different *Trypanosoma cruzi* strains treated with DETC. The parasites death cell mechanism was evaluated by flow cytometer as to the capacity for Annexin/Pi stain after 24 hours of treatment with different DETC concentrations; 4.44 µM; 44.44 µM; and 444.00 µM.

A second possible death cell mechanism was evaluated by the mitochondrial activity assay using rhodamine-123. For this, the parasites were treated with different concentrations of DETC (44.4 µM; 111.0 µM; and 222.0 µM) and, subsequently, the fluorescence rhodamine-123 stain was detected by flow cytometry. These concentrations were used to visualize the effect of DETC towards the parasite exposed for 24 hours, if 444.0 µM was used, the culture would be dead and it would not be possible to make inferences about experiment, since the stain was not present.

The fluorescent profile of rhodamine-123 stain in each strain of *T. cruzi* when expose to DETC is shown in Figure 5.A, the results was quantified and showed in Figure 5.B. Is possible observe that, in all strains, the exposure and increase of DETC concentration results in reduction of fluorescence emitted by parasites when compared with fluorescence emitted by parasite untreated. The strains Y (1) and QMM9 (4) were the least susceptible strains, with a similar profile of reduction in fluorescence emission, and the Dm28c (2) strain was the most susceptible to DETC, resulting in smaller fluorescence compared with control.

**Figure 4:**
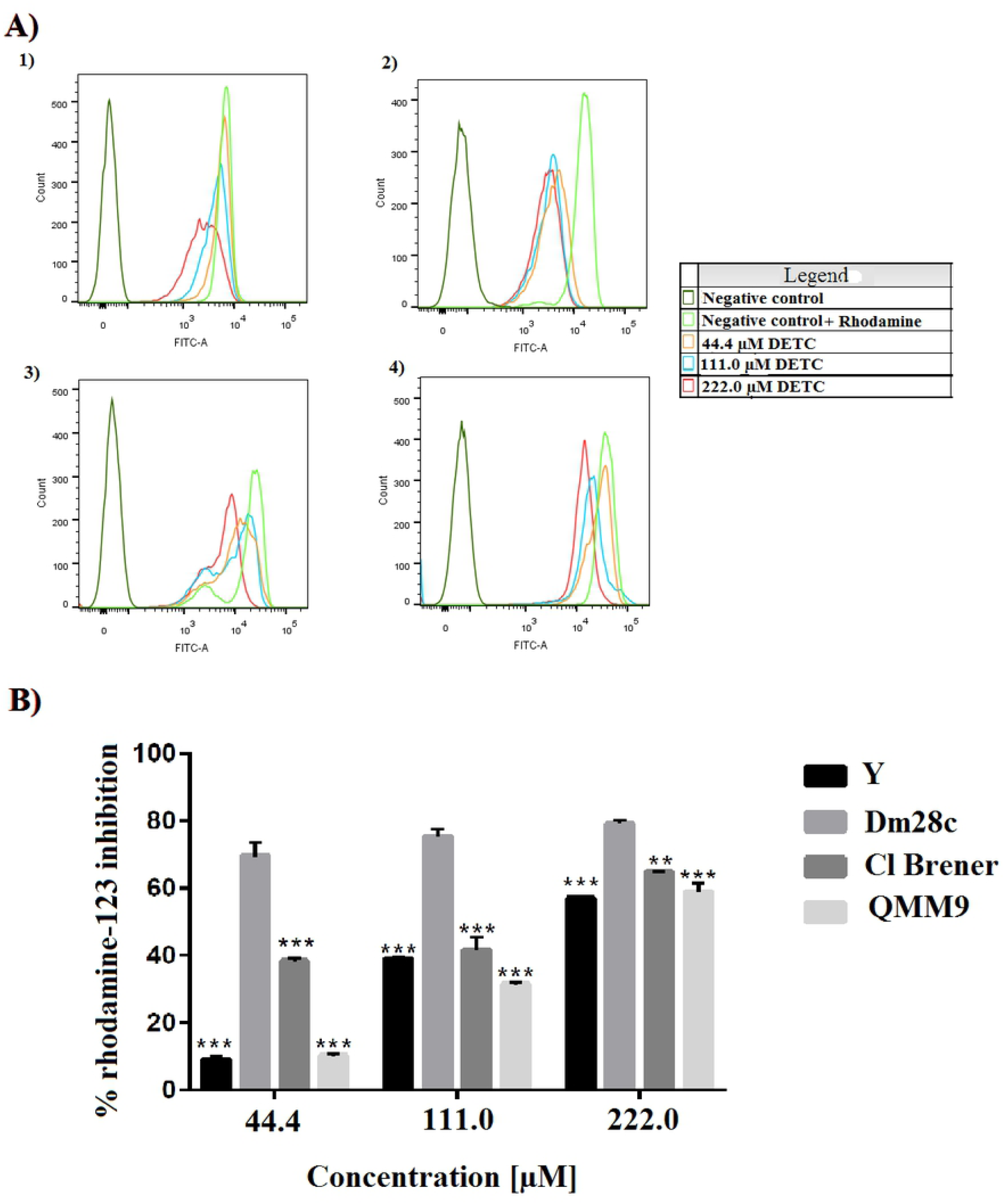
Evaluation of the mechanism of mitochondrial damage caused by DETC in *Trypanosoma cruzi*. A: Fluorescence intensity (Rhodamine-123) in different strains in trypomastigote form of *T. cruzi* untreated (control), represents 100% of fluorescence, and treated with different concentrations of DETC (44.4 µM, 111.0 µM, 222.0 µM) for 24 hours. (1) Y strain; (2) Dm28c strain; (3) Cl Brener strain; (4) Qmm9 strain. 10,000 events were used for each analysis to obtain the data, which was carried out in triplicate. B: Percentage of rhodamine-123 stain inhibition of different strains of *T. cruzi* treated with different concentrations of DETC. Results presented as mean ± standard deviation of the percentage of parasitic inhibition in a triplicate system and for the statistical analysis the Anova Test together with the Tukey Post-test (P <0.01 (**); P <0.001 (***) were used to compare the response in each concentration by strains towards different DETC concentrations.

## 4. Discussion

Chagas disease affects millions of people in endemic areas of Latin America and in several other non-endemic countries due to migratory flows of populations to countries in Europe, the United States, among others. In this scenario, this neglected tropical disease represents an important public health problem, especially in developing countries [31].

The parasite responsible for Chagas disease is the flagellated protozoan called *T. cruzi*. Although the etiologic agent is made up of only one species, *T. cruzi* has a genomic diversity that plays an important role in the epidemiological scenario of Chagas disease. The seven main genotypes of *T. cruzi* (TcI to TcVI) and Tc Bat are involved in the different epidemiological scenarios of the disease, mainly about pathogenesis, geographical distribution, reservoir and different degrees of susceptibility to pharmacological treatment [7-8].

Given the importance of the genetic diversity of *T. cruzi* in the context of the pharmacological treatment of Chagas disease, the objective of this work is to evaluate the antiparasitic activity of DETC *in vitro* against four different genotypes of *T. cruzi*, namely Dm28c strain (TcI), Y strain (TcII), QMM9 strain (TcIII), and Cl Brener strain (TcVI).

In the context, here we demonstrate the antiparasitic activity of DETC in the four different strains of *T. cruzi*. We observe different levels of resistance and susceptibility among these four different strains (Figure 1; Table 1). Regarding the evolutionary forms of the parasite, we also observed different resistance and susceptibility profiles of the epimastigotes and trypomastigotes of *T. cruzi* to DETC. The processes of *T. cruzi* metacyclogenesis are directly associated with changes in gene expression patterns, thus justifying a different profile of susceptibility to DETC [7-8,32-33].

The phylogenetic and evolutionary factors of the parasite also seem to be important variables when we analyse the antiparasitic activity of DETC against *T. cruzi*. In this study we demonstrated that DETC had lower IC_50_ values against non-hybrid strains of *T. cruzi*, such as Dm28c (TcI) and Y (TcII). However, highest IC_50_ values were obtained in the hybrid strains, QMM9 (TcIII) and Cl Brener (TcVI). Thus, the hybridity phenomenon in *T. cruzi* seems to have an important role in the rational design of new drugs for the control of Chagas disease. Evolutionary events can carry genetic characteristics of precursors that make them less susceptible to DETC, thus requiring greater concentrations to eliminate them [34-35]. In addition, hybridization processes are directly related to the development of strains resistant to certain drugs [36-37]. However, the mechanisms associated with resistance of different strains are yet unknown.

The antiparasitic activity of DETC against different strains of *T. cruzi*, presented in this work, is probably due to the chemical structure of this molecule, since the amino portion of DETC acting in reactive oxygen species production, thus justifying the possible mechanism of antiparasitic activity of this molecule [38]. In addition, the disulfide carbon group, present in the DETC molecule, could contribute to the interaction with metal centres present in numerous parasite enzymes [39].

DETC was identified as a group with activity against tripanosomatide family. In studies with *Leishmania amazonensis*, this compound showed the capacity of inhibition metallpreateases of parasite, in this case, iron superoxide dismutase in low concentrations of the compound, this caused an increment of ROS inside parasite leading to death [20-21]. In addition, *in vitro* studies have shown that DETC have the capacity to inhibit antioxidant enzymes of *T. cruzi*, increasing radical superoxide and radical peroxide inside parasite [39-40]. These mechanisms can be associated with superoxide inhibition, an important enzyme of redox metabolism and essential to parasite survive [41-42].

The data presented in Figure 2 contribute to the fact that oxidative damage could lead the parasites to death. The presence of pores and membrane disruption after treatment with DETC, were observed by other authors after treatment with compounds that cause oxidative damage [43-45]. Therefore, we believe that oxidative damage produced by DETC can trigger morphological changes in *T. cruzi*, causing structural changes that result in the death of the parasite.

With regard to the DETC cytotoxic profile, in this study we observed the viability of the 3T3 and RAW cell lines, treated with DETC for 24 hours (Figure 3). Only in high concentrations, DETC was able to, significantly, reduce the viability of these cells, but not less than 40%. Even at high concentrations, DETC was not toxic enough to cause 100% mortality of cell lines. However, even in low concentrations, DETC has 100% antiparasitic activity.. With these data it was also possible to determine the selectivity index (SI) of DETC of different strains and evolutionary forms of *T. cruzi* (Table 2). The results presented in Table 2 show the same profile of the BNZ SI compared to the 3T3 cell line when compared to the fibroblast cell line [46]. Thus, DETC has less toxicity than BNZ when evaluated in RAW cell lines [47]. These results corroborated the evaluation of *in vitro* efficacy of DETC compared with conventional drug used in CD treatment.

Understanding possible mechanisms of action of the anti-parasitic activity of DETC against *T. cruzi* was also one of the objectives of this work. We noticed that the results observed with annexin/PI stain were contrary when compared with results present in Figure 1, 1S and 2. As it was expected a high stain was associated with cellular process of death, but in Figure 4, an absence of stain in strains Dm28c and Y were observed, and a smaller stain in QMM9 and Cl Brener. Initially, we believed that the kit was spoiled, we tested with two new kits, but the results gave the same output (results not shown). In addition, we believe that DETC could be spoiled, and we retested the resazurin essay, but the results were similar, present in figure 1, so it confirmed the DETC efficiency. At the same time, we evaluated the antiparasitic activity of DETC against *Leishmania amazonensis*, where it can be seen that DETC at 44.4 µM is capable of marking 50% of parasites by the annexin/PI assay (results not shown).Other possibility was that the protocol provided by kits was not adequate to *T. cruzi*. However, your laboratory recently published a study using the same *T. cruzi* treated with silver nanoparticles (100 µg/mL) and annexin/pi stain detected was 100% [48]. We do not know why *T. cruzi* treated with DETC did not marked annexin/PI. We speculate that this phenomenon is due to the ability of DETC to induce anti-apoptotic pathways in the parasite, acting as a regulator of apoptosis, since the ability of DETC to act on caspases through the thiol group of the molecule, consequently inhibiting signal transmission to trigger apoptotic processes [49-50]. This mechanism was observed in some kind of cell lines after treatment with DETC [51-54]. On other hand the difference of stain between strains could be associated with phylogenetic processes. However, these speculations needs to be further studied and characterized.

In this work, we also observed that increasing the concentration of DETC decreases both the metabolic capacity of resazurin and rhodamine-123, showed on Figure 1 and 5. It is known that the metabolism of these two compounds by the parasite is directly related to the functioning of complexes I, II and III of the inner membrane of the mitochondria [55]. The phenomenon of reduction of these compounds may be related to the damage of the inner membrane of the parasites’ mitochondria, resulting from the presence of DETC and causing a malfunction in the regulation of the protonation gradient carried out by complexes I, II and III and impairing ATP synthesis, and consequently leading to parasite death [55-58]. Interestingly, it can be seen in Figure 5 that the highest concentration of DETC (222.0 µM) is able to reduce the metabolism of rhodamine-123 in all strains of *T. cruzi* analysed in this study, probably associated with the reduction of the membrane potential mitochondrial, potentiating the mitochondria as a possible target for the anti-*T*.*cruzi* activity presented by DETC.

The cytoplasmic membrane damage observed in *T. cruzi* treated with DETC occasioning the formation of pores can also be directly related to the damage of the parasite’s mitochondria, because DETC has the ability to cause mitochondrial damage. Some studies report the capacity of mitochondrial damage caused by the production of reactive species, a possible explanation for the antiparasitic mechanism presented by the DETC [59-61].

In conclusion, in this work we demonstrated the antiparasitic activity of DETC against different evolutionary forms (epimastigotes and trypomastigotes) and four different strains of *T. cruzi*, the etiological agent of Chagas disease. When we compared the antiparasitic activity of DETC with BZN, the first-line drug for the treatment of Chagas disease, DETC showed better antiparasitic activity *in vitro*. Finally, we believe in the need for new pharmacological tests in animal models in order to enhance the use of DETC as a promising candidate in the treatment of Chagas disease.

## Acknowledgments

We would like to thank Prof. João Aristeu da Rosa and Dr. Aline Rimodi Rimeiro at UNESP Araraquara (Brazil) for offering four different strains of *T. cruzi*. JWFO, CJGM, and BAC thanks to the financial support (PhD and Post-doctoral fellowships) provided by Capes/Brazil; MSS and HAOR thanks to CNPq/Brazil for the Research Grant (*Bolsa de Produtividade*). We also would like to thank the Department of Materials Engineering at UFRN for allowing the use of their scanning electron microscope, and the Department of Biochemistry at UFRN for allowing the use of their Flow Cytometer. We are also grateful to Paulo Fanado for editing this manuscript.

## Author contributions

Experimental study design: JWFO, TNT, HAOR and MSS; Cultivation of parasites and assessment of antiparasitic activity: JWFO, CJGM, MSS; Flow cytometer assays and data analysis: JWFO, AKMCS, JSB, HAR; Scanning electron microscopy methodology and data analysis: JWFO, BAC, and IZD; Obtaining research funding: HAR and MSS. In addition, all authors actively participated in the writing and discussion of the manuscript. All authors read and approved the final version of the manuscript.

## Supporting Information

**Figure 1S.**
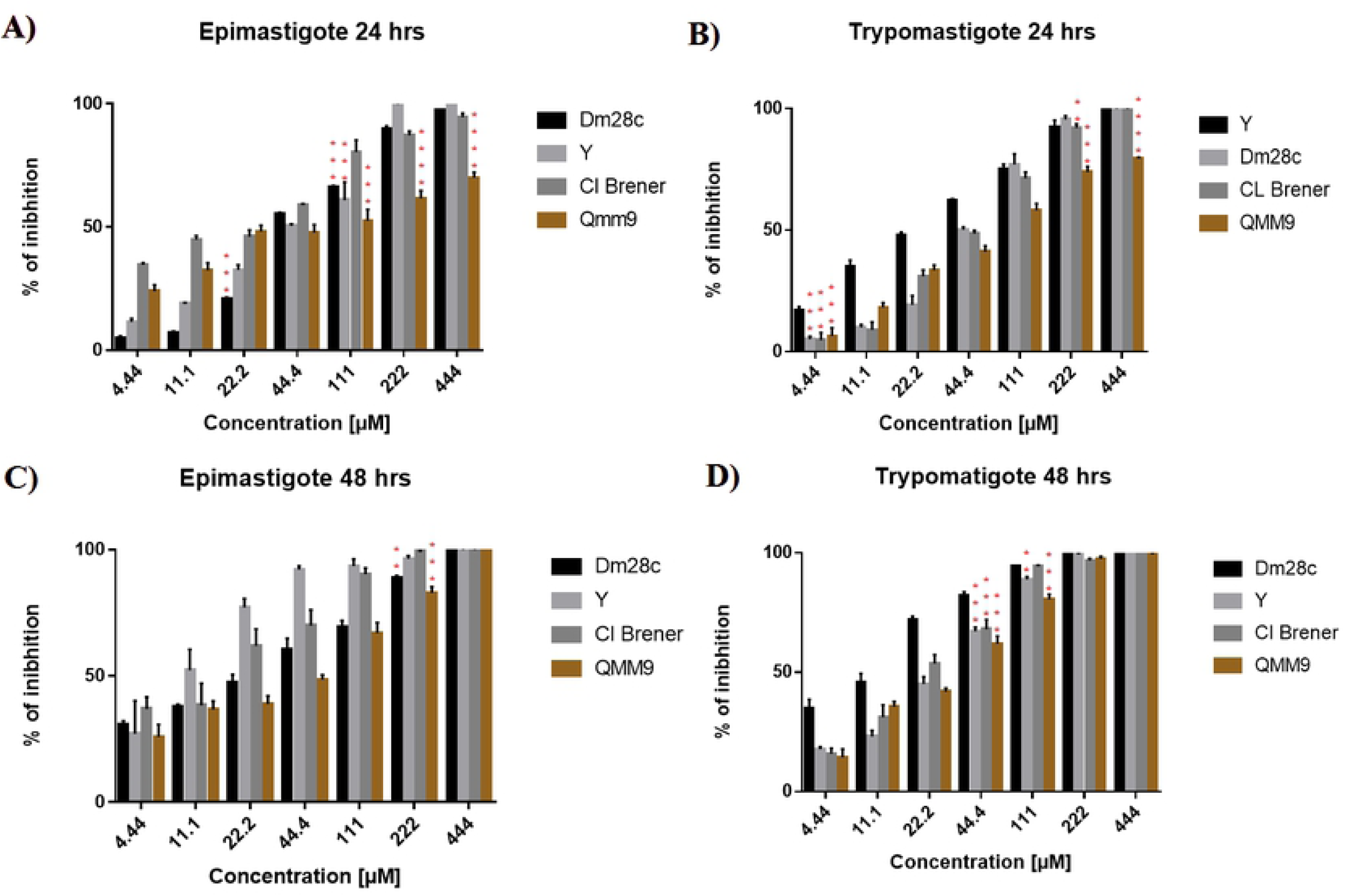
Percentual of inibhition in different strains of *T. cruzi* after treatment with DETC with range concentration 4.44 µM until 444.0 µM analysed by count. A) Epimastigote 24 hours, B) Trypomastigote 24 hours, C) Epimastigote 48 hours and D) Trypomastigote 48 hours. Results presented as mean ± standard deviation of the percentage of parasitic inhibition in a triplicate system and for the statistical analysis the Anova Test together with the Tukey Post-test (P <0.01 (**); P <0.001 (***), P<0.0001(****). In statistical analysis was compared the response in different strains submitted to same concentration of DETC.

